# Different behaviourally relevant stimuli evoke different forms of adaptation in the olfactory system

**DOI:** 10.1101/2024.09.12.612693

**Authors:** Elliot Birkett, Anton Nikolaev

## Abstract

Sensory neurons are continuously exposed to a large diversity of stimuli and adjust their response according to the recent stimulation history, in a process called adaptation. Recent studies have demonstrated that many sensory neurons not only depress (decrease their response to repetitive sensory stimulation) but some neurons exhibit facilitation: a small initial response followed by increase in response amplitude. Adaptation has been mainly studied with neutral stimuli and it is not known whether different behaviourally relevant stimuli evoke adaptation with similar or different properties. Here we used ethologically relevant stimuli to study adaptation in the zebrafish olfactory system. We found that repetitive presentation of food odour caused a variety of adaptations ranging from very strong depression to facilitation, but depression was the predominant dynamic. On the other hand, a different behaviourally-relevant stimulus, alarm substance, evoked much stronger facilitation and also decreased the amplitude of responses to food presentation. Other sensory modalities, aversive mechanosensory and attractive visual stimuli, also caused adaptation with different properties in the same brain area. Different forms of adaptation may therefore be used for processing sensory stimuli evoking different behavioural reactions.

**Graphical abstract:** 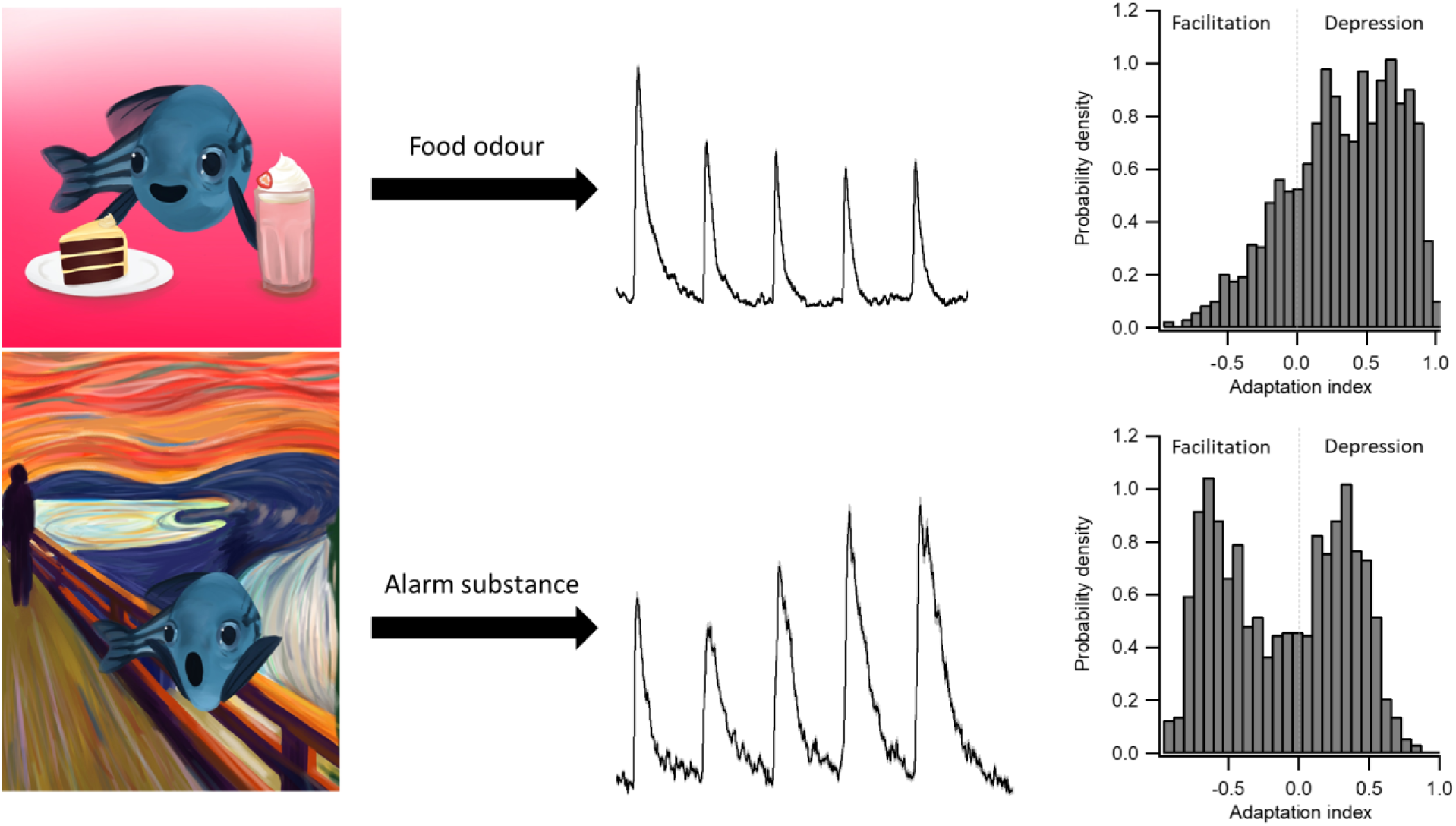

**Highlights:** Food odour evokes a diversity of adaptations ranging from extreme depression to facilitation

Alarm substance evokes much stronger facilitation than food odour

Alarm substance inhibits responses to food odour

Mechanosensory stimulation evokes both depression and facilitation

Visual stimuli evoke responses in the olfactory bulbs that strongly and quickly depress

## 1. Introduction

The brain has to perform complex computation while being exposed to a large number of sensory stimuli that signal the presence of food, danger, conspecifics and neutral (irrelevant) objects (Colwill et al., 2005; Kermen et al., 2013; Privat et al., 2019). Upon receiving information from different sensory modalities, the brain has to ignore neutral objects, and execute specific behavioural programs to stimuli that are relevant to animal’s survival and/or reproduction (Angelaki et al., 2009; Del Bene et al., 2010; Mu et al., 2019). When dealing with a large number of diverse stimuli, sensory neurons adjust their response amplitude depending on recent stimulus history. This can be expressed in two different forms: adaptation, a change of response amplitude to repetitive presentations of the same stimulus; and cross-stimulus modulation, a change of response amplitude to the same stimulus in the presence of another stimulus.

Adaptation arises at the early stages of sensory systems (e.g. in the retina, hair cells or olfactory bulbs) and has multiple mechanisms. Visual adaptation in retinal ganglion cells, for example, results from the intrinsic inactivation of sodium channels (Kohn, 2007), reduced glutamate release through synaptic depression at bipolar cell terminals or direct inhibition onto retinal ganglion cells or bipolar cells via amacrine cells (Baden, 2017; Demb, 2008; Manookin and Demb, 2006; Nikolaev et al., 2013; Rieke and Rudd, 2009). Recent studies in mammalian, amphibian and zebrafish visual systems have found a rich diversity of adaptation dynamics(Kastner and Baccus, 2011; Nikolaev *et al*., 2013).

Following increases in visual contrast, bipolar and ganglion cell synapses with strong initial responses depress, while those with initially weak responses facilitate (Kastner and Baccus, 2011; Nikolaev *et al*., 2013). This pattern was anticorrelated with the inhibitory feedback from amacrine cells. Pharmacological block of GABA receptors led to a switch of facilitating synapses to depressing ones, suggesting that facilitation results from depression of the inhibitory feedback from amacrine cells. Together, depression and facilitation allow for efficient transfer of sensory information, with depressing neurons able to respond accurately to increase in stimulus strength and facilitating cells able to respond accurately to decreases from the input system. Such opposing forms of plasticity have been found in other visual areas (Heintz et al., 2022) as well as the mechanosensory system, (Pichler and Lagnado, 2019) but it is not known whether they exist in all sensory systems.

Despite the fact that mechanisms of sensory adaptation are largely known, the function of adaptation is still unclear. It is considered to be important for sharpening the acuity of stimulus detection, removing redundancy and modulating the stimulus salience, but these possibilities have not been decisively proven (Kohn, 2007; Solomon and Kohn, 2014). So far, the adaptation was mainly studied using neutral stimuli such as white noise or full field light (Nikolaev *et al*., 2013) to stimulate the visual system, or amino-acids (Jacobson et al., 2018) to stimulate the olfactory system. These stimuli have little ethological (behavioural) relevance and it is not known whether sensory stimuli executing different behaviours cause adaptation with different dynamics. Probing adaptation with ethologically relevant stimuli may clarify its role of adaptation for sensory processing (Solomon and Kohn, 2014).

Here, we studied the adaptation of neuronal responses in the olfactory bulbs and ventral telencephalon (Dp) to two types of odours (food and the alarm substance, Schreckstoff), as well as visual and mechanosensory stimuli. Both olfactory stimuli evoke different and strong behavioural reactions – attractive (food) and aversive (Schreckstoff) (Jesuthasan et al., 2020). We show that food odour triggers a diversity of adaptations ranging from extreme depression to facilitation but the majority of the neurons exhibited depression.

On the other hand, the alarm substance evoked less depression and more facilitation, as well as cross-odour modulation – the amplitudes of neural responses to the food odour were strongly inhibited by the alarm substance. To explain these results, we propose that in order to process the information about food, the brain needs to perform a variety of computations (novelty detection, stimulus recognition, stimulus localisation, etc.) where different forms of adaptation may be beneficial (Kastner and Baccus, 2013; Solomon and Kohn, 2014). In contrast, the alarm substance, which triggers predator avoidance behaviour, requires fast processing. Here, an increase to the sensitivity to the alarm substance would signal persistent danger. Contrasting adaptation to stimuli of different ethological relevance was further supported by a different set of sensory stimuli, an attractive visual stimulus and repulsive mechanosensory stimulation, which evoked depression and a mix of depression and facilitation, respectively.

## Materials and Methods

### Animals and Husbandry

All experimental procedures were performed according to UK Home Office regulations (Animals (Scientific Procedures) Act 1986) and approved by the Ethical Review Committee at the University of Sheffield.

Zebrafish were bred through either standard pair mating or breeding over marbles procedures (see https://www.sheffield.ac.uk/bateson/zebrafish/information). Embryos were then raised in E3 solution without methylene blue (5 mM NaCl, 0.17 mM KCl, 0.33 mM CaCl_2_, 0.33 mM MgSO_4_). Larvae were maintained at 28°C and housed on a 14/10 light/dark cycle. After 5 days post fertilisation (dpf), larvae were transferred to a techniplast system (Techniplast S.p.A., Buguggiate, Italy) where they were fed twice a day, excludings the day of experiment.

### Imaging and Stimulation Application

Multiphoton calcium imaging of zebrafish neurons was conducted using the pan-neuronal *tg*(*Xla.Tubb:GCaMP3*) (Bergmann et al., 2018) and *tg*(*elavl3:GCaMP6s*) (Chen et al., 2013). Non-anaesthetised 4-5dpf larvae were immobilised in 3.5% low melting point prepared with E3 solution prior to imaging. Subjects were mounted dorsal side up onto a custom built chamber (Fig.1A). Older 10-17 dpf larvae were mounted in an oxygenated 3.5% agarose solution as described in (Bergmann *et al*., 2018). In both cases, the agarose surrounding the olfactory region was removed to aid olfactory detection.

**Fig.1.**
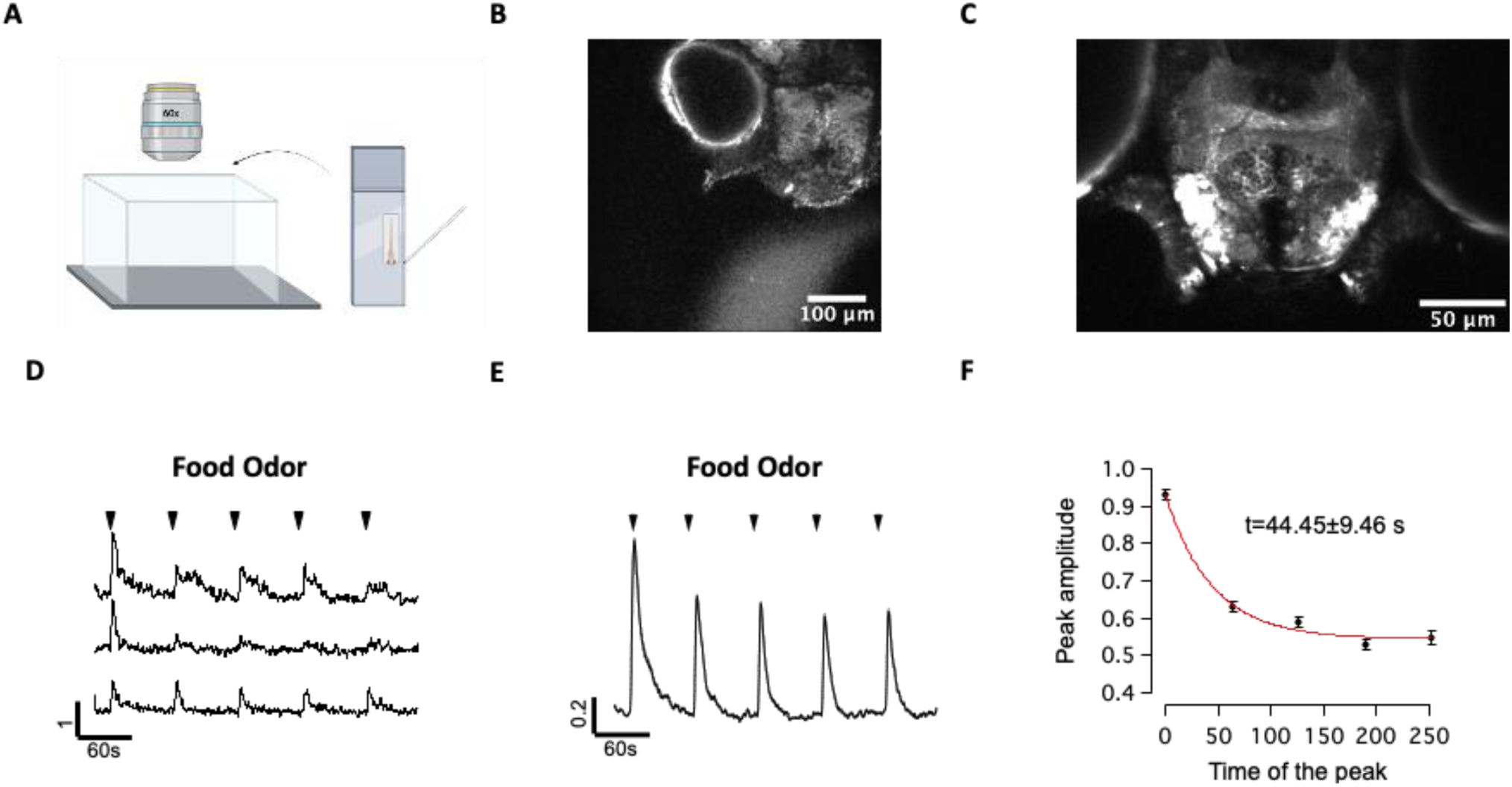
Zebrafish olfactory system exhibits adaptation to repetitive application of food odour. **A,B** - Imaging set up. Fish was immobilised in 3% agarose with the area around the nose freed. Odours were delivered as puffs via pipette in a way so that the stream is not directly oriented towards the nose. This was tested in a separate set of experiments where fluorescent dye was included in the odour delivering pipette (B). Note that such a way of stimulation still caused some mechanosensory stimulation (Fig. 5A). **C** - Example of an imaging area with olfactory bulbs and the Dp area. **D** - Example of fluorescence dynamics of individual voxels **E** - Average fluorescence dynamics from 1750 voxels in response to repetitive presentation of 5 odour puffs. SEM is shown in grey. On average, repetitive application of food odour causes depression. **F** - Average adaptation can be fitted with a single exponent with a time constant of 44.45±9.46s, which is slightly slower than in the visual system.

Olfactory stimuli were delivered using a picospritzer set to 3 psi for a duration of 3 seconds. Stimulus delivery was achieved through a 10 μm diameter glass pipette. The stimuli presented consisted of an E3 control, a food-based stimulus, and the alarm substance, schreckstoff. The food-based stimulus and alarm substance were obtained as described in (Kermen et al., 2020), and (Jesuthasan *et al*., 2020), respectively. All solutions were filtered to remove all physical contaminants and applied anterior to the nose of the fish to minimise mechanosensory stimulation.

Imaging was conducted on a commercially available two-photon laser-scanning microscope (Bergamo II System, Thorlabs Inc., USA) based on a mode-locked laser system operating at 925 nm, 80-MHz pulse repetition rate, < 100-fs pulse width (Mai Tai HP DeepSee, Spectra-Physics, USA). Images were captured using a 60× objective, 1.1 NA (LUMFLN60XW, Olympus, Japan) using a GaAsp PMT (Hamamatsu) coupled with a 525/40 bandpass filter (FF02-525/40-25, Semrock) as described in (De Faveri F., 2021). Single plane responses were recorded at 15 frames per second at a resolution of 512 x 512 pixels.

Experimentation began with a 30-second base line fluorescence recording before the first presentation of the odorant was applied. A total of 5 presentations of any given olfactory stimulus were provided with a 30 second inter trial interval.

Visual stimuli were generated using custom written code for Matlab (MathWorks, Natick, MA, USA) with the Psychophysics Toolbox using an Optoma PK320 projector connected to a Linux laptop. Stimuli were projected onto the side of a custom built fish cinema that passed across the visual field of the left eye (dimensions restricted to - 90 mm (W) × 15 mm (H) due to working distance of the objective). The eye-to-screen distance was 1.3cm.

### Data Analysis

Raw data files obtained from 2-photon imaging were registered using TurboReg (Bergmann *et al*., 2018) for ImageJ to remove motion artefacts and every 5 frames were averaged to reduce noise artefacts. The forebrain was then split through the midline so analysis could be conducted separately for the left and right side of the brain and improve ease of data handling.

Analyses were performed using SARFIA (Dorostkar et al.,2010) and custom-written scripts for Igor Pro (WaveMetrics, Lake Oswego, OR, USA) as described in (Bergmann *et al*., 2018). For manual sorting, analysis was performed on a voxel-wise basis. Fluorescence intensity was normalised and individual voxels selected as responding by calculating the skewness of the distribution curve. Traces with a skewness > 1.0 were regarded as responding based on work described in (Bergmann *et al*., 2018). To aid with sorting, all responding traces were re-organised based on their similarity to one another based on the similarity index (Pearson coefficient) and then every 5^th^ trace sorted into categories based on their adaptive properties.

Additional analysis was also performed to avoid potential bias from manual sorting. Registered files were once again uploaded to Igor Pro where a region of interest (ROI) detection plugin (Johnston J, 2019) identified regions displaying similar patterns of calcium activity. Voxels within each ROI were averaged to provide a single trace displaying the fluorescence activity in this area. ROIs were then sorted based on their adaptation index defined as below:

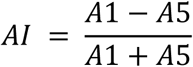

A histogram of AI values was used to identify thresholds of depressing, sensitising and non-adapting traces.

### Statistical Analysis

Statistical analysis was performed using Igor Pro version 6.3. All average traces are shown as mean ± S.E.M. Differences in adaptation index between response types and before and after PTZ application were analysed using two-tailed Wilcoxon rank-sum test. Data before and after drug application were considered independent. The non-normal distribution of adaptation indices is shown and validated through the Jarque-Bera test. Data collection was not performed blind to the conditions of the experiment.

## Results

To understand whether different behaviourally relevant stimuli evoke adaptation with different properties in the same brain area, we used a set of sensory stimuli including two olfactory stimuli (odour of food and odour of alarm substance, Figs. 1-4), one visual stimulus (small spot moving horizontally, Fig. 4B) and a mechanosensory stimulus (a water jet of E3 solution without any odour included, Fig. 4A). Each of these stimuli has been previously shown to evoke distinct behavioural reactions: presentation of food odour or a small spot triggers appetitive/hunting behaviour (Bianco et al., 2011), alarm substance evokes freezing behaviour in larval zebrafish (Jesuthasan *et al*., 2020), and mechanosensory stimuli elicit escape responses (Eaton et al., 2001). Using this diverse set of behaviourally relevant stimuli we aimed to define potential differences in adaptation dynamics of neuronal activity in the olfactory bulbs and ventral telencephalon.

### Depression and facilitation in the olfactory system

Food and alarm substance odours were delivered 5 times for a duration of 3 seconds with 60 second interstimulus intervals. Both olfactory stimuli were delivered in standard E3 solution via a glass pipette, without directly targeting the nasal area. This was confirmed by including a fluorescent dye (Lucifer Yellow, Fig. 1A, B) in the patch pipette during a control experiment. We used this approach as an alternative to standard constant application of a stream jet directed at the nose because the latter causes some mechanical stimulation during the odour change and because constant mechanosensory stimulation evokes adaptation (De Faveri F., 2021; Pichler and Lagnado, 2019). It would therefore make comparison of adaptation to olfactory and mechanosensory stimuli difficult. However, our results suggest that neither depression nor facilitation in the olfactory system can be attributed to mechanosensory stimulation alone (see below).

Analysis was performed on the early olfactory system including olfactory bulbs, ventral telencephalon (Fig. 1C) and the olfactory epithelium (Fig. 4A). We performed voxel-wise analysis in which fluorescence dynamics of individual voxels were analysed separately without defining whether they belong to the same cell. However, similar results were also obtained when analysis was performed on the soma of individual neurons (not shown).

Repetitive presentation of the food odour caused a significant depression – decrease of the response amplitude (Fig. 1D, E). The average amplitude of the first response was about 1.7x higher than that of the fifth response (0.96 ± 0.016 vs 0.55 ± 0.017, p<0.0001, n = 1755 voxels, 5 fish, Fig. 1F), giving an average adaptation index of 0.27±0.01. Adaptation dynamics can be fitted with a single exponent, with the adaptation time constant (tau) being 44.45 ± 9.46 seconds, which is slower than in the visual system (2-10 seconds, (Nikolaev *et al*., 2013)). Thus, zebrafish olfactory neurons exhibit strong adaptation, which is in agreement with previous studies(Friedrich and Laurent, 2004).

Visual systems of mammals and other vertebrates exhibit different forms of adaptation with varying degree (depressing vs facilitating) and speed of adaptation (Baccus, 2012; Kastner and Baccus, 2013; Meister, 2002; Nikolaev *et al*., 2013). Therefore, we aimed to define whether olfactory neurons also exhibit such diversity of adaptations. To test this, we categorised voxels based on the adaptation dynamics of their fluorescence and averaged the response traces (Fig.2). Responding traces were categorised into three main subgroups: depressing (34.7%, adaptation index = 0.65 ± 0.01), non-adapting (18.9%, adaptation index = −0.04 ± 0.001) and facilitating (15.9%, adaptation index = - 0.24± 0.01). Although most of the traces exhibited an increase in fluorescence upon application of food odour, some also behaved like OFF cells, i.e. decreased fluorescence upon food odour application and/or subsequently depolarized at the stimulus offset (Supplementary Fig. 1). This data demonstrates that, like in the visual system, the olfactory system exhibits opposing forms of plasticity in response to food odour presentation.

**Fig.2.**
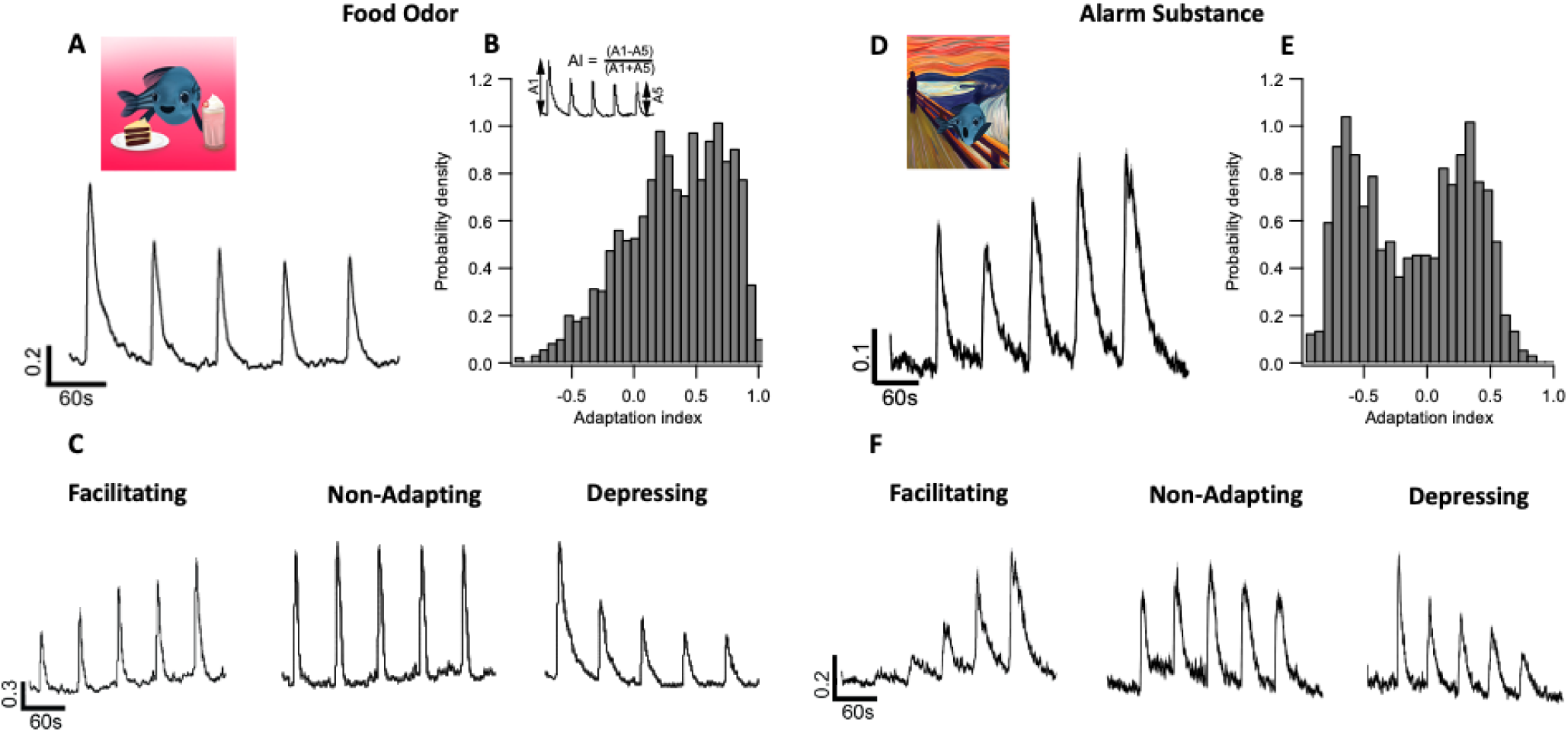
Alarm substance (Schreckstoff) causes much stronger facilitation than food odour. **A,D** - Average responses to 5 presentations of food odour or alarm substance respectively. Notice the difference in the degree of adaptation. **B**, **E** - Distribution of the adaptation index in response to food odour (left) and Schreckstoff (right). Insert on B explains how adaptation index was calculated - it was defined as a ratio of difference and sum of the first and fifth response amplitude. Note the bimodal nature of the alarm substance distribution, highlighting two types of adaptation - facilitation (AI< 0) and depression (AI>0). Notice the larger number of facilitating voxels in response to alarm substance. **C, F** - Different types of responses to repetitive presentation of food or alarm substance odours. Voxels were manually sorted and categorised into 3 clusters for each odourant: depressing, non-adapting and facilitating. An additional cluster (OFF cells) was also observed (Supplementary Fig.1) in response to food odour. The voxels were averaged for each cluster and plotted to demonstrate the different adaptation dynamics. Error bars for all plots shown in grey.

The experiments above were conducted on young larval zebrafish (<5dpf). The observed diversity of adaptations could, therefore, be attributed to the immature brain or to stimulus novelty - larval fish will not have experienced this olfactory stimulus before. Consequently, we investigated whether the diversity of adaptation responses was preserved during development by investigating the adaptation of olfactory neurons at later larval stages (10-17dpf, Supplementary Fig.2). We found a similar representation of adaptation as described for the 5dpf zebrafish, suggesting that opposing forms of plasticity are not a temporary characteristic of the undeveloped olfactory system and are not limited to stimuli that were not experienced by the animal before.

### Food and alarm substance evoke adaptation with different properties

The experiments above demonstrate that food odour evokes a range of adaptations in the zebrafish olfactory system. As an attractive stimulus, the presence of food odour drives a set of behavioural programmes, such as appetitive and exploratory behaviour (Kermen *et al*., 2020). Will other sensory stimuli that evoke other behaviours (e.g. aversive sensory stimuli) cause adaptation? We hypothesised that because such stimuli signal persistent danger, they should evoke less depression and more facilitation. To test this, we first compared adaptation dynamics to two olfactory stimuli on the same fish (attractive food odour and repulsive alarm substance, Fig. 2-3) and then compared these to adaptation caused by other sensory modalities(mechanosensory and visual) in the same brain areas (Fig.5).

**Fig.3.**
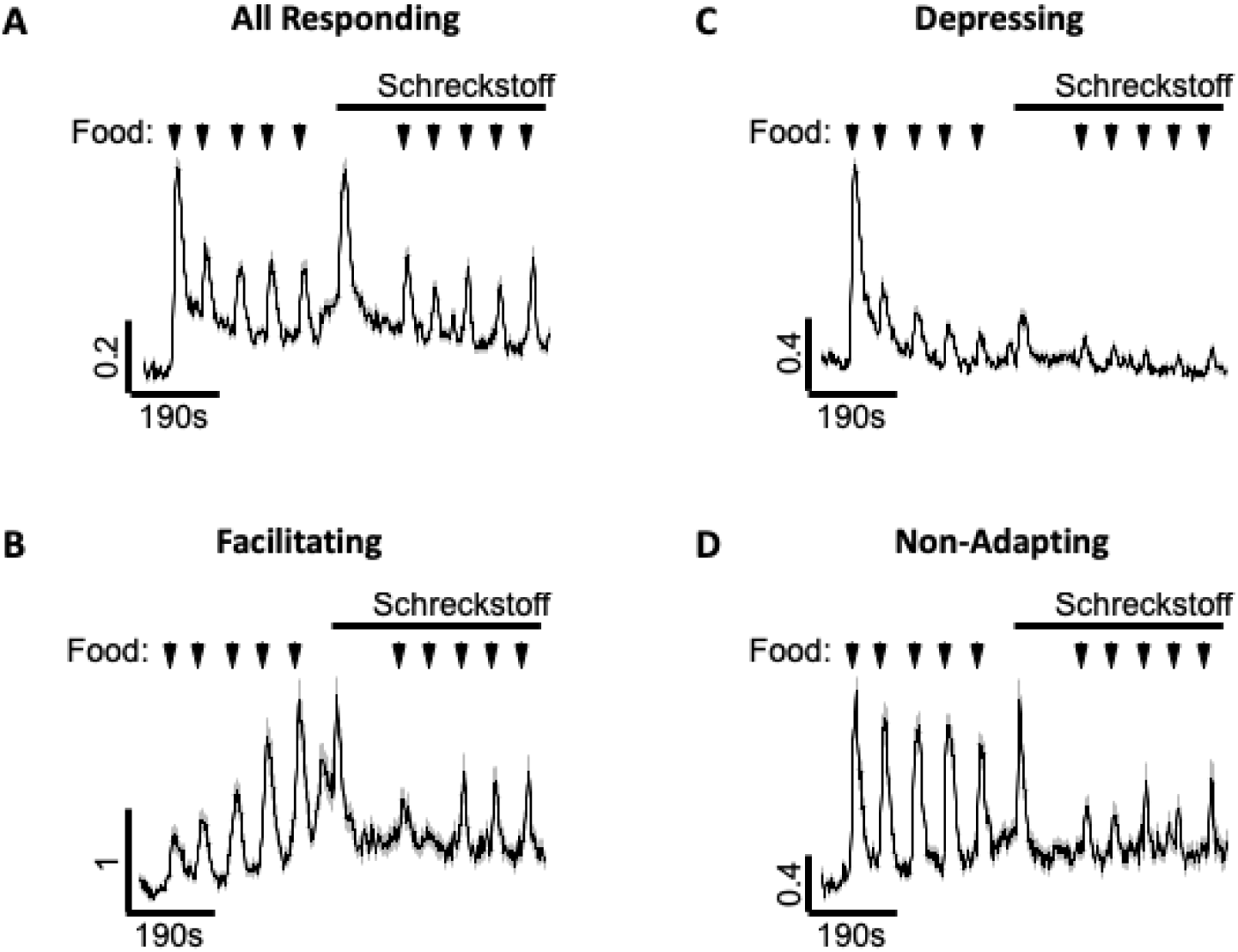
Schreckstoff inhibits responses to food odour. **A** - Average experiments where food odour was applied 5 times (arrows), followed by prolonged application of Schreckstoff and subsequent additional applications of food odour (second set of 5 black arrows). **B**-**D** - Responses of facilitating(**B**), depressing (**C**) and non-adapting (**D**). Note that facilitating and depressing terminals show much smaller responses to food odour after Schreckstoff application and therefore decrease in response to food after Schreckstoff application cannot be explained by depression in response to food odour.

On average, food and alarm substance cause adaptation with different dynamics. While food caused depression (average AI = 0.29 ± 0.01, n = 1755), the alarm substance evoked some facilitation (Fig. 2A, D, average AI = −0.16 ± 0.01, n = 1245, p<0.05, nonparametric two-sample Wilcoxon test).

We then compared the adaptation dynamics in food- and alarm substance-evoked adaptation at the level of individual voxels. In both cases we found a diversity of adaptation dynamics, ranging from depression to facilitation (Fig.2, C, F). Distribution of AI, especially in alarm substance, revealed two peaks (AI = - 0.68 and AI= 0.29), displaying both depression and facilitation in roughly equal proportions (Fig2, B, E). However, food mostly evoked depression with the majority of the traces exhibiting AI>0. Therefore, we conclude that although both odours evoke a diversity of adaptation dynamics, food evokes mainly depression while alarm substance evokes both depression and facilitation.

#### Error bars shown in grey

The above results reveal some aspects that are important for processing of olfactory stimuli. The adaptation to our olfactory stimuli was not complete. Even to our food stimulus that evoked stronger depression, there were always some neurons within the olfactory bulbs still responding to sensory stimulation. This ensures some information regarding the odorant was able to reach higher processing centres. Alarm substance on the other hand signals immediate danger and threat. It is therefore a perfect processing strategy to not decrease, and even increase the sensitivity to a stimulus that repeatedly signals the presence of danger. However, other sensory signals could occur that may put the fish at further risk to predation should it trigger exploratory behaviours and override the initial freezing response to the alarm substance. We therefore hypothesised that the presence of the alarm substance may cause zebrafish to stop responding to other stimuli and thus assist with the freezing behaviour evoked by the alarm substance (Jesuthasan *et al*., 2020).

To test this, five applications of food were applied anterior to the nose of the fish, followed by an application of an alarm substance and then a further five applications of the food stimulus (Fig. 3A). On average, responses were seen to all 10 presentations of food odorant, suggesting the olfactory system is still sensitive to other stimuli after exposure to the alarm substance (Fig. 3A). However, the average response amplitude to the food odorant was reduced after alarm substance exposure (p<0.005, n = 934).

Because the whole population of neurons depresses on average, the above result can be explained by depression of responses to food odour rather than inhibition of response to food by alarm substance application. To rule this out, we have analysed the effect of alarm substance in each adapting class (Fig. 3B-D) and found that the effect of alarm substance was observed in facilitating and non-adapting terminals and therefore cannot be explained by depression of food responses alone (p < 0.05, n = 44). Taken together, these results demonstrate that the alarm substance inhibits the responses of olfactory neurons to food odours.

### Different mechanisms of facilitation in the visual and olfactory system

The above results indicate that the olfactory system, like visual and the lateral line systems (Baccus, 2012; De Faveri F., 2021; Nikolaev *et al*., 2013; Pichler and Lagnado, 2019), exhibit both depression and facilitation. In the visual system the main mechanism of facilitation results from the depression of the inhibitory feedback to early visual neurons (Kastner and Baccus, 2013; Nikolaev *et al*., 2013). However, in the olfactory system, the mechanism of facilitation appears to be different: facilitation was quite prominent in the olfactory epithelium (Fig.4A) with the majority of the olfactory neurons showing prominent facilitation in response to the alarm substance. This is important because olfactory epithelium neurons do not receive any inhibitory feedback either locally or from downstream areas, and therefore, should have different mechanisms of facilitation.

**Fig.4.**
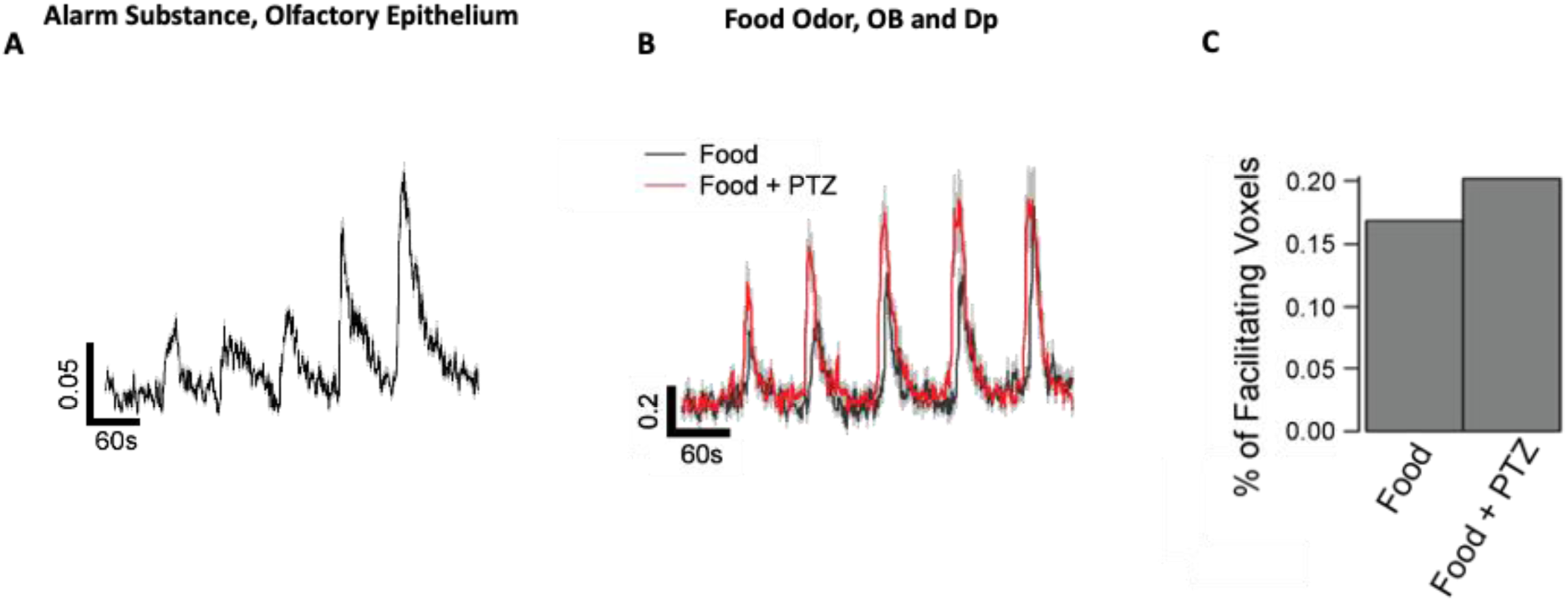
Olfactory system appears to have different mechanisms of facilitation than visual system. **A** - Response to Schreckstoff in the olfactory epithelium. Notice profound facilitation despite the absence of inhibitory input to these neurons. **B**-**C** - Inhibition of the negative feedback does not turn facilitation into depression. **B** - Adaptation dynamics before and after application of PTZ (inhibitor of GABA receptors) in the same fish. Notice that there is still facilitation after application of PTZ (red trace). **C** - percentage of facilitating voxels before and after PTZ application.

To test this idea further, we have compared the facilitation dynamics to food application in absence and presence of GABA_A_ receptor inhibitor pentylenetetrazole (PTZ) in the same animal, thus blocking inhibitory feedback. Fig.4B shows responses of facilitating voxels in absence (black) and presence of 100 mM PTZ. The overall response amplitude was higher and the terminals still show strong facilitation in presence of PTZ. Importantly, the proportion of the facilitating terminals did not decrease (Fig.4C, 18% vs 20%). Taken together, these two experiments demonstrate that the mechanisms of facilitation in the olfactory system do not result from depression of a local inhibitory feedback.

### Adaptation to other sensory modalities

#### Error bars shown in grey

The results above indicate that different behaviourally relevant stimuli evoke adaptation with different dynamics in the olfactory bulbs. Our attractive stimulus, food odour, evoked mainly depressing responses, while the alarm substance which signals immediate danger evokes a mix of depression and facilitation. It therefore appears that in the fish subpallium, the attractive stimuli cause more depression, while repulsive stimuli can evoke more facilitation. We have further tested this idea with a different set of sensory stimuli from other modalities - an attractive visual stimulus, a small moving spot resembling food and a repulsive mechanosensory stimulation by puff of E3 solutions. These stimuli evoke strong behavioural reactions (Bianco *et al*., 2011) and it was shown that both of them activate the sensory neurons in the ventral telencephalon(Shainer et al., 2023).

Application of E3 solution evoked responses in the olfactory bulb and the ventral telencephalon and repetitive presentation of E3 evoked adaptation with different properties (Fig.5A and Supplementary Fig.3). Both depression and facilitation were observed in response to E3 and application of food odour evoked stronger depression than E3 alone.

**Fig.5.**
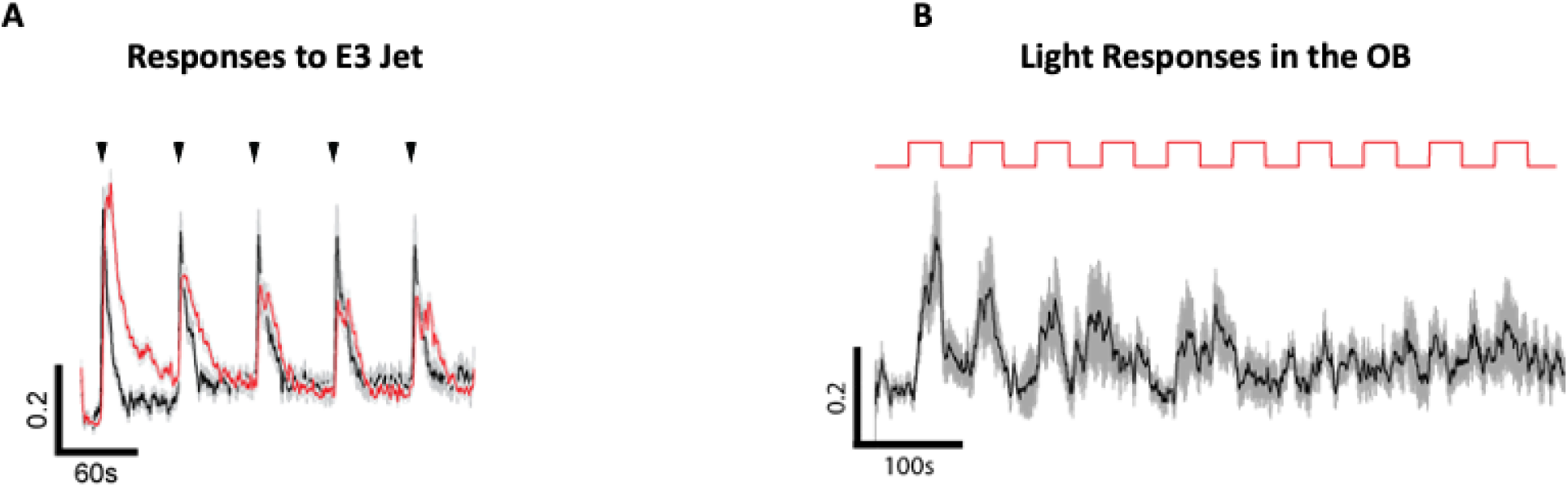
Average responses of olfactory bulbs and Dp area to light and mechanosensory stimulation. **A** - Response to 5 puffs of E3 solution without any odour (black) and food odour in the same fish (red). Notice that on average, responses to E3 exhibit much less adaptation. **B** - Responses to repetitive visual stimulation in the olfactory bulbs - small black spot moving horizontally across the visual field and projected to a screen. Notice that the adaptation is quick and complete.

The number of depressing, non-adapting and facilitating terminals was much higher in response to food odour than E3 alone (330 vs 132, all responding, 128 vs 38, depressing and 41 vs 28 facilitating. N = 13 fish). This is particularly drastic in the olfactory epithelium, where E3 alone did not evoke any response, while application of the alarm substance evoked strong responses with strong facilitation (Fig.4A). We therefore conclude that both olfactory and mechanosensory stimuli evoke diverse forms of adaptation.

To evoke light responses we used a dark spot that triggers approaching behaviour when moving horizontally (Bianco *et al*., 2011). On average, this caused very quick depression (Fig.5B) with the majority of voxels responding to the first presentation only.

## Discussion

In this paper, we show that the olfactory system of zebrafish exhibits depression and facilitation at the level of olfactory sensory neurons, olfactory bulbs and the ventral telencephalon. Facilitation in these regions is more pronounced to two repulsive sensory stimuli (alarm substance and mechanosensory stimulation) whilst depression is more prevalent in the presence of attractive stimuli (smell of food and small moving spot). In addition, we also show that alarm substance has the capacity to inhibit olfactory responses to other stimuli - food odour. This phenomenon will likely improve the animals ability to survive during dangerous situations through preventing unwanted behavioural exploration.

Probing olfactory systems, particularly in water is challenging because it always activates some mechanosensory system. Because the lateral line shows some degree of facilitation, the facilitation in response to odours could actually be explained by facilitation in response to mechanosensory stimulation. However, this cannot be the case. First, the number of facilitating neurons depends on the type of olfactory odourant (Fig. 2B vs Fig. 2E). Second, the number of facilitating terminals in the same fish is much smaller in response to E3 vs food. Third, facilitation is observed at the level of olfactory epithelium but there is very little response there to just E3 mechanosensory stimuli. We conclude that facilitation to olfactory stimuli is genuine.

### Mechanisms of facilitation in the olfactory system

We show here that like in other sensory systems, the neurons of the olfactory system adapt in different ways - some depress while others facilitate. The mechanisms of facilitation in the olfactory system differ from those observed in the visual system and cannot be explained by depression of the inhibitory feedback(Nikolaev *et al*., 2013). This is because they occur in the olfactory epithelium that lacks inhibitory cells and also because pentylenetetrazol, the inhibitor of GABA_A_ receptors, has a minimal effect on adaptation dynamics. This raises the question: what are the mechanisms of facilitation in the olfactory system? While the topic is a subject of further studies, two potential mechanisms can be proposed:

1. Sensitivity of olfactory receptors increases during response to alarm substance. This mechanism seems unlikely as it requires a complex response of a number of receptors encoding a specific odour.
2. Increase in overall excitability of olfactory receptors during stimulation due to increase in the activity of voltage-gated sodium channels or decrease in the activity of potassium channels. Testing this hypothesis would require patch clamping individual olfactory receptors.

### Function of facilitation

The function of adaptation, particularly facilitation, remains a subject of debate. Initially, facilitation in sensory systems was thought to primarily function to improve the responses to decreases in stimulus intensity(Kastner and Baccus, 2013; Nikolaev *et al*., 2013). However, this role might be better suited to contrast-suppressed cells or OFF cells. In the retina, these cells can signal a sudden decrease in light luminance and contrast. Notably, OFF cells were also found in our experiments in the olfactory bulbs. They can also be suitable to signal sudden decreases of the odour concentration (Supplementary Fig. 1) due to fish movement and therefore assist in chemosensory motor behaviour (Daghfous et al., 2012). Therefore, the original explanation for the function of facilitation may not be entirely complete.

Another proposed function for facilitation is the preprocessing or “whitening” of sensory data, particularly in the visual system where low frequencies tend to depress and high frequencies tend to facilitate. Yet, this role does not apply to olfaction.

Before our research, adaptation had not been compared with behaviorally relevant stimuli. Our findings reveal that certain stimuli evoke depression, while others evoke facilitation. A pattern begins to emerge: substances like alarm substance and water jets signal danger, while food and small spots signal nourishment. Thus, the function of facilitation might be to increase the weight of neurons responding to dangerous signals and by that enhancing the organism’s ability to respond to threats.

## Acknowledgments

We thank members of Jesuthasan and Marcotti labs for comments on the manuscript. This work was sponsored by the Royal Society and BBSRC grants (AN) and A* Star PhD studentship (EB).

## SUPPLEMENTARY FIGURES

**Supplementary Fig.1.**
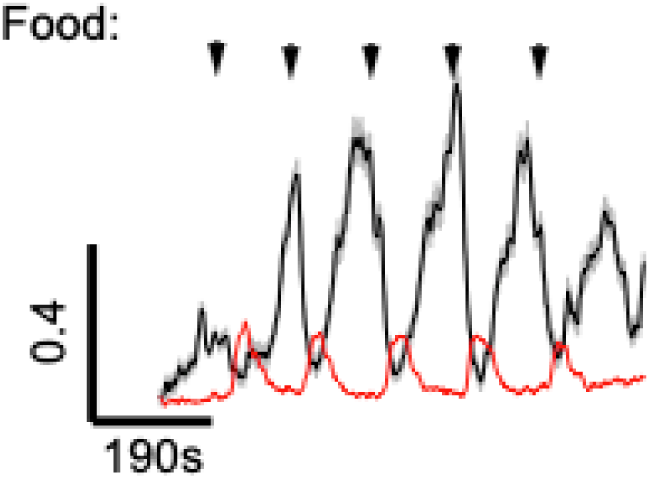
Although most of the cells responded to food odour by activation, some neurons responded by inhibition. Average response of OFF cells is shown in black. Red - average response to food odour shown for comparison. See also supplementary Fig. 2 for another example from 17dpf fish.

**Supplementary Fig.2.**
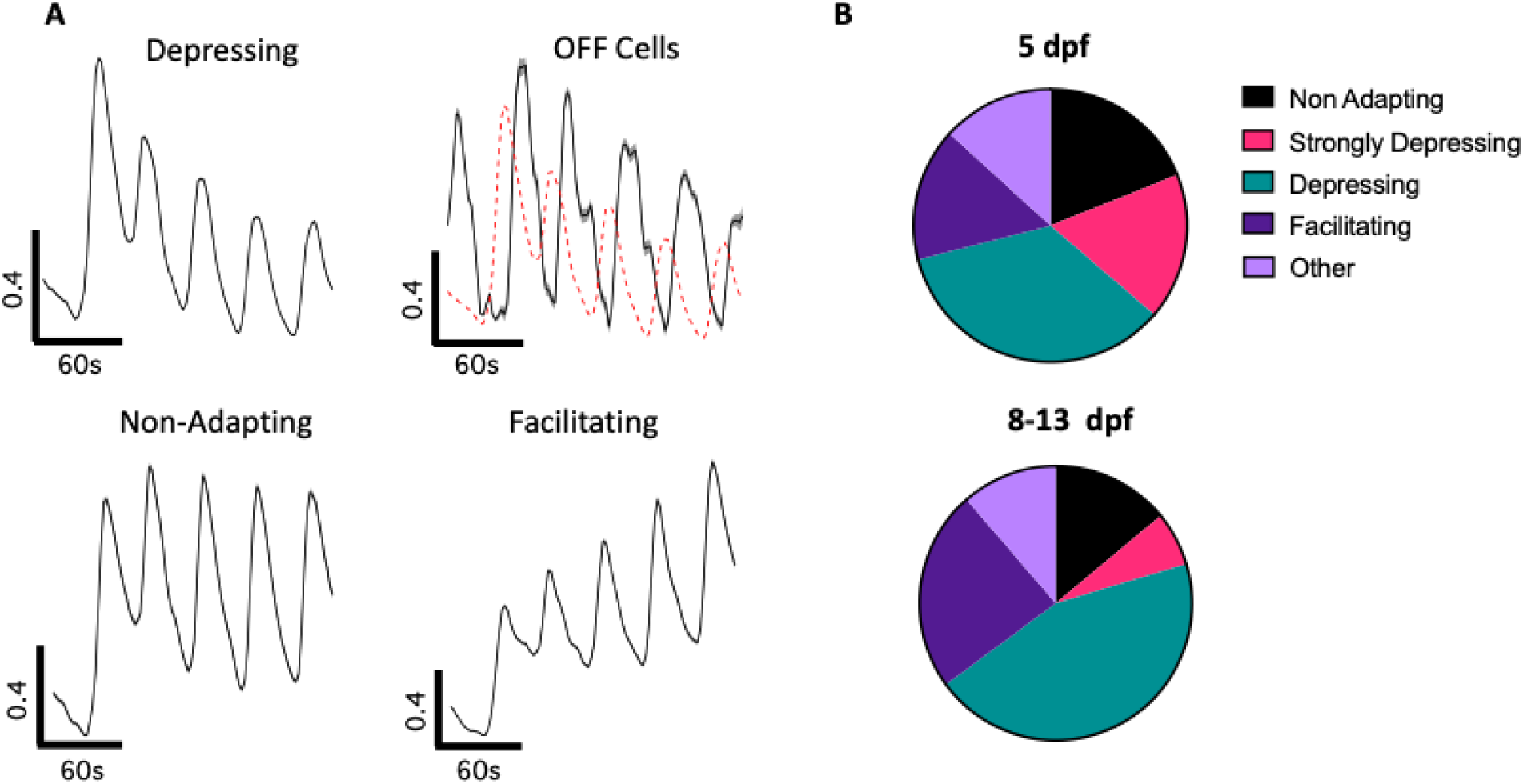
Different dynamics of adaptation are observed in late stage larvae. **A** - Different types of responses, including depressing, non-adapting, facilitating and OFF in 10-17 dpf larvae. Responses were analysed using single voxel analysis and clustered using cluster analysis using SARFIA (Dorostkar et al., 2010). **B** - Relative distribution of each cluster at 5 and 17 dpf. Distribution is similar, with the largest change simply in the degree of depression. This suggests that diversity of adaptations in the olfactory system is not due to immaturity of the olfactory system.

**Supplementary Fig.3.**
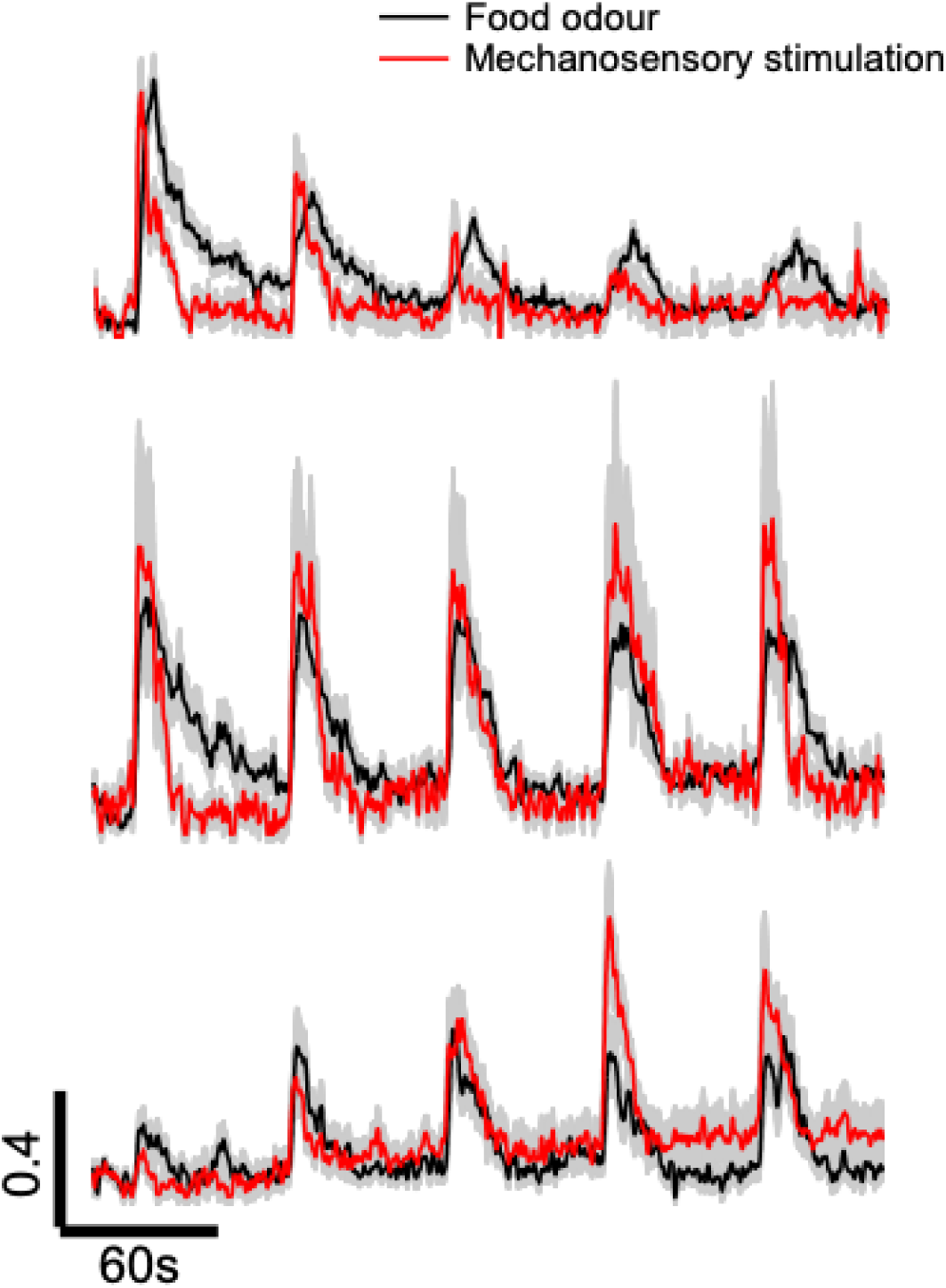
E3 mechanosensory stimulation evoked facilitation and depression in the olfactory bulbs. Different types of adaptation dynamics were seen in response to both food odour and mechanosensory stimulation in the same fish.

## Notes

### Competing Interest Statement

The authors have declared no competing interest.

